# Genetic regulation of the bacterial omega-3 polyunsaturated fatty acid biosynthesis pathway

**DOI:** 10.1101/2020.01.28.924217

**Authors:** Marco N. Allemann, Eric E. Allen

## Abstract

A characteristic among many marine Gammaproteobacteria is the biosynthesis and incorporation of omega-3 polyunsaturated fatty acids into membrane phospholipids. Biosynthesis of eicosapentaenoic (EPA) and/or docosahexaenoic (DHA) acids is accomplished using a polyketide/fatty acid synthase mechanism encoded by a set of five genes *pfaABCDE.* This unique fatty acid synthesis (FAS) pathway co-exists with the canonical Type II dissociated fatty acid synthesis pathway, which is responsible for the biosynthesis of saturated, monounsaturated, and hydroxylated fatty acids used in phospholipid and lipid A biosynthesis. In this work, a genetic approach was undertaken to elucidate genetic regulation of the *pfa* genes in the model marine bacterium *Photobacterium profundum* SS9. Using a reporter gene fusion, we showed that expression of the *pfa* operon is down regulated in response to exogenous fatty acids, particularly long chain monounsaturated fatty acids. This regulation occurs independently of the canonical fatty acid regulators, FabR and FadR, present in *P. profundum* SS9. Transposon mutagenesis and screening of a library of mutants identified a novel transcriptional regulator, which we have designated *pfaF*, to be responsible for the observed regulation of the *pfa* operon in *P. profundum* SS9. Gel mobility shift and DNase I footprinting assays confirmed that PfaF binds the *pfaA* promoter and identified the PfaF binding site.

**Importance:** The production of polyunsaturated fatty acids (PUFA) by marine Gammaproteobacteria, particularly those from deep-sea environments, has been known for decades. These unique fatty acids are produced by a polyketide-type mechanism and subsequently incorporated into the phospholipid membrane. While much research has focused on the biosynthesis genes, their products and the phylogenetic distribution of these gene clusters, no prior studies have detailed the genetic regulation of this pathway. This study describes how this pathway is regulated under various culture conditions and has identified and characterized a fatty acid responsive transcriptional regulator specific to the PUFA biosynthesis pathway.

## Introduction

Regulation of fatty acid biosynthesis, particularly the levels of unsaturated fatty acids, has been shown to be a crucial aspect of the bacterial physiological response to a variety of environmental conditions, including temperature, pH, and hydrostatic pressure. Both biochemical and transcriptional regulatory mechanisms exist to regulate the various aspects of the fatty acid biosynthetic pathway (1). In the model organism *Escherichia coli*, genes that compromise the Type II fatty acid synthase (FAS), and in particular genes related to monounsaturated fatty acid (MUFA) biosynthesis are regulated by the interplay between FadR and FabR (2–4) as shown in Figure 1. FadR is a member of the GntR regulator family and it acts as positive regulator of both *fabA* and *fabB*, which encode for proteins essential to the biosynthesis of unsaturated fatty acids in this organism (5). As seen in Figure 1 exogenous fatty acids (>C10) are imported across the outer membrane via the FadL transporter and subsequently converted into acyl-CoAs by the acyl-CoA synthetase FadD. In the absence of long chain acyl-CoA, FadR binds to sites upstream of *fabA* and *fabB* and acts as a positive regulator (6, 7). Upon acyl-CoA binding to FadR, it adopts a conformation that is unable to bind to its cognate sites upstream of *fabA/B* promoters leading to deactivation of transcription. FabR, a TetR family transcriptional regulator, further regulates *fabA/B* by acting as a classical repressor. FabR binds to sites immediately downstream of the FadR site in both *fabA/B* regardless of acyl-CoA being present (3, 8). While an exact role of FabR has yet to be described, it has been speculated that the opposing actions of FabR and FadR at *fabA* and *fabB* promoters is ultimately responsible for the regulation of these genes (3).

**Figure 1.**
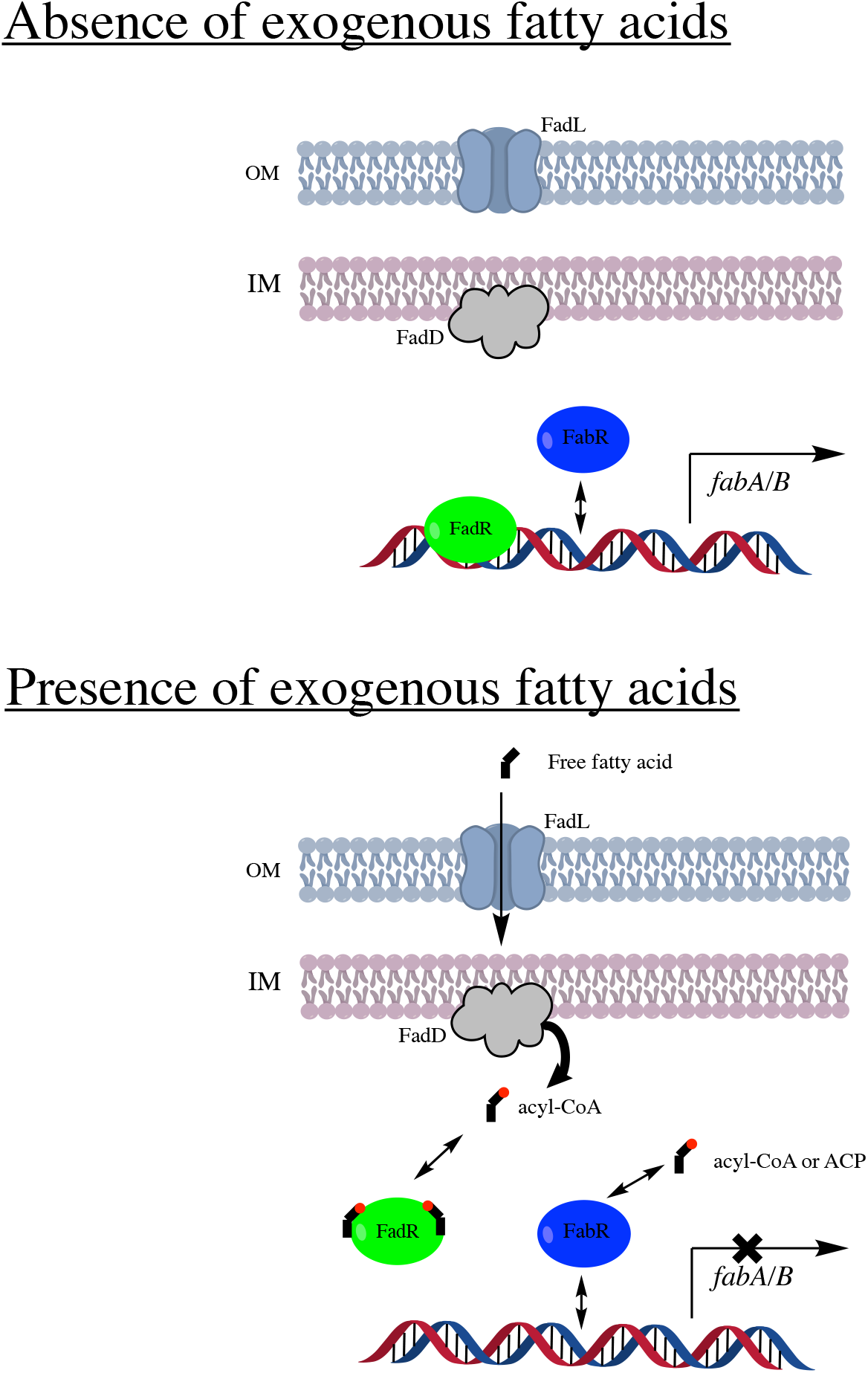
Genetic regulation of monounsaturated fatty acid biosynthesis genes *fabA* and *fabB* in *E. coli*. In the absence of exogenous fatty acids FadR binds to a site upstream of the *fabA* and *fabB* promoters and acts as an activator of transcription. When present, exogenous fatty acids are transported across the outer membrane by FadL and converted into acyl-CoA by FadD. Acyl-CoA binding to FadR causes a conformational shift that abolishes the DNA binding capabilities of FadR. In both scenarios FabR binds to a site downstream of FadR and has been shown to bind in the presence/absence of acyl-CoA and/or Acyl-ACP. Loss of FadR activation of transcription presumably allows FabR to act as a better repressor of *fabA/B* expression.

Similar regulatory mechanisms regulating monounsaturated fatty acid biosynthesis have been characterized in other model Gram-negative bacterial systems. In *Shewanella oneidensis* MR-1 the FabR homolog was found to be responsible for the regulation of *fabA* and *desA*, an oxygen dependent membrane bound lipid desaturase (9). Similar regulatory mechanisms controlling unsaturated lipid biosynthesis have been described in other model bacteria such as *Pseudomonas aeruginosa* PAO1, which lacks a *fadR* homolog (1, 10, 11). In this strain *fabA* and *fabB* form an operon (unlike *E. coli*), which is regulated by another TetR family regulator, DesT (12). In addition to *fabAB* expression, DesT also modulates the expression of the membrane bound desaturase DesBC, that catalyzes oxygen dependent desaturation of saturated acyl-CoA which can then be incorporated into membrane phospholipids (10, 11).

A subset of marine Gammaproteobacteria, particularly strains isolated from cold and/or high-pressure environments, produce omega-3 polyunsaturated fatty acids (PUFA) such as EPA (20:5*n-3*) and DHA (22:6*n-3*), that are incorporated into phospholipid membranes (13, 14). The biosynthesis of these unique fatty acids is linked to the *pfaABCDE* operon, which encodes for a type I FAS/polyketide synthase (15). In these bacteria, the Pfa synthase pathway co-exists with the Type II FAS, which produces saturated (SFA) and monounsaturated fatty acids (14, 16). Given that both pathways utilize the same precursor substrates (15, 17, 18) and their respective end products are destined for phospholipids (13, 14, 19–22), an interesting question arises as to how these pathways are physiologically coordinated in the cell. Numerous studies have demonstrated that culturing native PUFA-producing strains at cold temperature (23–28) and/or high pressure (23, 26) leads to increases in PUFA abundance. In the EPA producing bacterium *Photobacterium profundum* SS9, analyses of transcript abundances of the *pfa* operon at cold temperatures and/or high pressure indicated no significant alterations mRNA abundances relative to 15°C or low-pressure conditions (25). In the course of those studies, a chemical mutant of *P. profundum* SS9 (EA2), was shown to have increased mRNA abundance relative to its parental strain indicating the possibility of a transcriptional regulator(s) existing in the strain (25).

In this work, the transcriptional regulation of the *pfa* operon in *P. profundum* SS9 is further characterized and shown that it is down regulated in response to exogenous unsaturated fatty acid (18:1) supplementation. A genetic screen utilizing a *pfaA∷lacZY* reporter gene fusion combined with transposon mutagenesis was used to identify a novel transcriptional regulator, herein designated *pfaF*, which positively regulates the *pfa* operon and mediates the regulatory response to exogenous fatty acids. Gel mobility shift and assays using purified PfaF demonstrated that PfaF binds to the *pfaA* promoter and that this binding is sensitive to the presence of oleoyl-CoA. DNase I footprinting was used to determine the binding site of PfaF within the *pfaA* promoter.

## Results

### Expression of the *pfa* operon under various culture conditions

To better understand the regulation of the *pfa* operon and to facilitate monitoring of gene expression, a reporter construct was designed to link expression of the *pfa* operon to the *lacZY* operon of *Escherichia coli.* This *pfaA∷lacZY* strain allows the *pfa* operon promoter to be monitored in single copy with all possible upstream regulatory sequences. As expected from previous work (25) in *P. profundum* SS9, loss of *pfaA* rendered the *pfaA∷lacZY* strain unable to produce EPA (data not shown). Assays for β-galactosidase activity indicated that in mid-log phase growth approximately 48 Miller units of activity were produced (Figure 2A). Further β-galactosidase assays under conditions shown previously to lead to increased EPA content, such as high hydrostatic pressure and low temperature are given in Figure 2A and indicated no changes in LacZ activity, confirming previous results from this strain (25). Sequence analysis of the promoter region from EA2, a previously isolated EPA overproducing strain with increased *pfa* operon transcript levels, also indicated no changes in the promoter sequence of strain EA2 relative to the wild-type.

**Figure 2.**
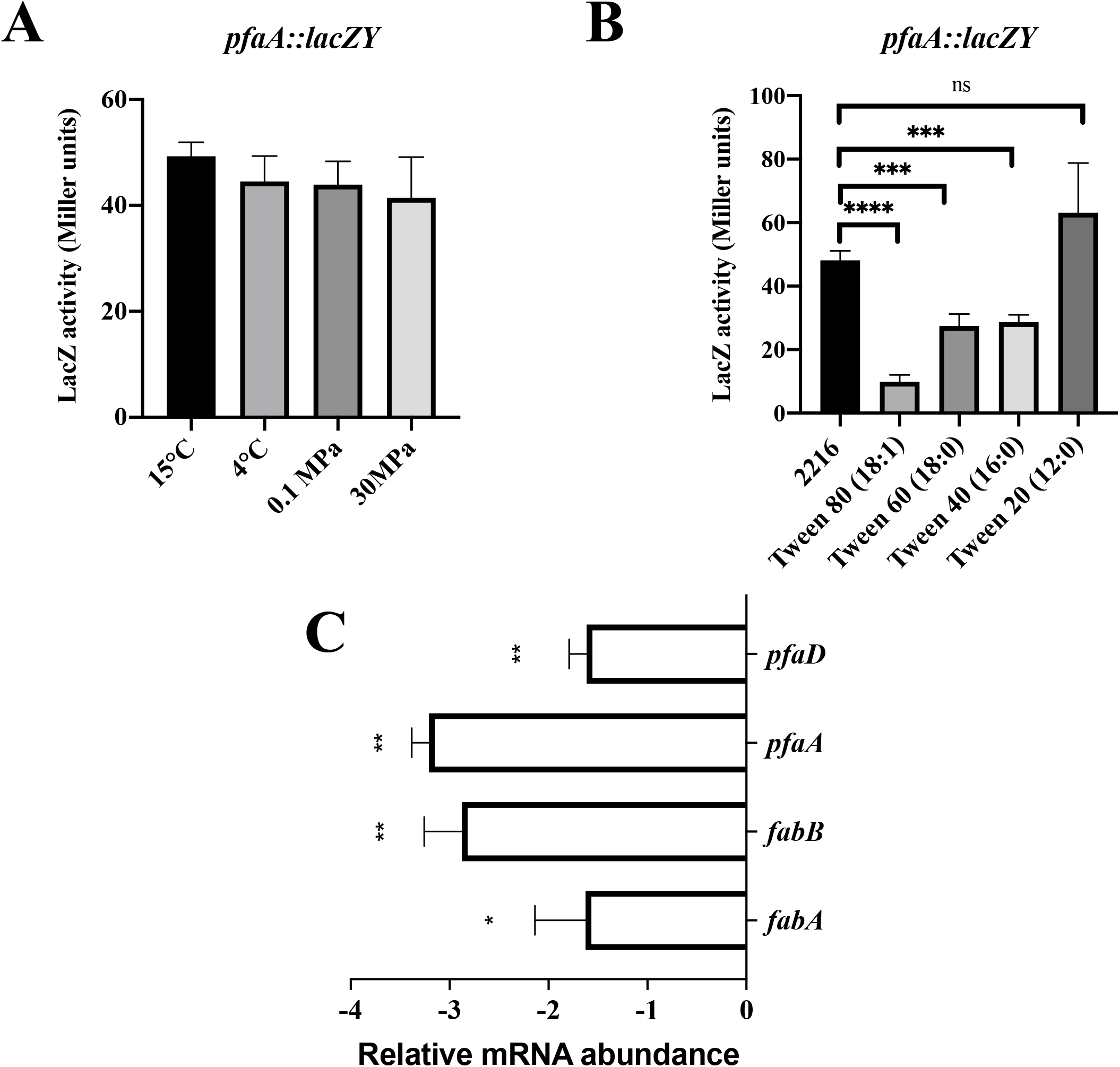
Reporter gene expression in the *pfaA*∷*lacZY* strain in response to a variety of culture parameters known to modulate EPA composition. (A) Various temperature and hydrostatic pressure conditions. (B) Cells cultured at 15°C with 0.05% Tween compound supplement indicated. For (A) and (B) results of at least six independent experiments shown as means with error bars representing standard deviation (ns, *P* >0.05; ***, *P* <0.005; ****, *P <*0.0001). (C) Effect of 0.05% Tween 80 supplementation on various fatty acid biosynthetic gene transcript abundances in SS9R as determined by qRT-PCR. Cells grown without supplementation represent the calibrator condition. Error bars represent the standard deviations based on at least three independent biological replicates with duplicate qPCR reactions (*, *P* <0.05; **, *P* <0.005).

Given its biosynthetic role in producing fatty acids destined for phospholipid biosynthesis, we hypothesized that the *pfa* operon might be regulated in a similar fashion to the prototypical *fab* regulon. Given the previously noted solubility issues of fatty acids in 2216 marine growth medium (23), exogenous fatty acids in the form of various polysorbate esters (Tween 20 (12:0), 40 (16:0), 60 (18:0), 80 (18:1)) were utilized as exogenous fatty acid supplements. As shown in Figure 2B, significant decreases in β-galactosidase activity were observed in the *pfaA∷lacZY* strain in response to all Tween compounds except for Tween 20 (12:0) (Figure 2B). Given the various degrees of down-regulation noted amongst the various Tween compounds seen in Figure 2B, which differ only in their fatty acid component, the possibility of this response being due to the polysorbate component of these compounds can be eliminated. This down-regulation was also shown to occur in SS9R as both *pfaA* and *pfaD* transcript abundances are reduced ~3-fold and ~2-fold, respectively, in response to Tween 80 (18:1) supplementation (Figure 2C). Transcript abundances of *fabA* and *fabB* were also decreased under these conditions (Figure 3C) indicating that, similar to the case in *E. coli* (2–4, 8), the monounsaturated fatty acid (MUFA) biosynthesis genes are down regulated in the presence of exogenous Tween 80 (18:1). Fatty acid profiling of SS9R grown in the presence of Tween 80 supplementation indicated a nearly 4-fold increase in 18:1 composition and an approximate 10-fold decrease in EPA consistent with the down-regulation of *pfaA* noted above (Supplemental Table 2). A novel fatty acid, tentatively identified as 18:2, was also observed in cultures supplemented with Tween 80 and may be representative of further metabolic processing of the incoming 18:1 acyl chain.

**Figure 3.**
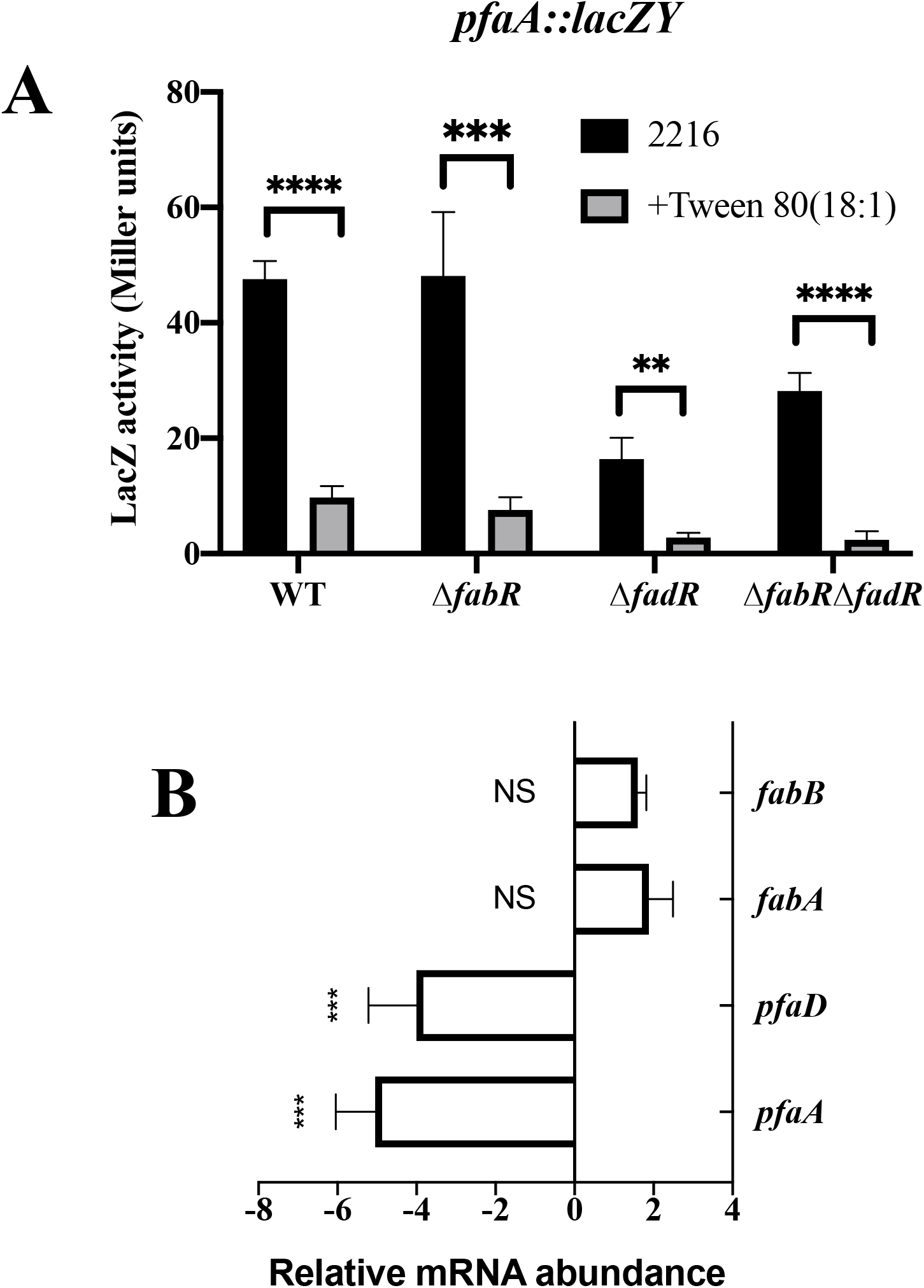
Influence of FadR and FabR on expression of the *pfa* operon. (A) LacZ activities of strains carrying Δ*fabR* and/or *ΔfadR* mutations in the presence or absence of 0.05% Tween 80 (18:1). Results of at least six independent experiments shown as means with error bars representing standard deviation. (**, *P* <0.05; ***, *P* <0.005; ****, *P <*0.0001) (B) Relative transcript abundances of *pfaA, pfaD, fabA*, and *fabB* grown in the presence or absence of Tween 80 (18:1) in (SS9R *ΔfabRΔfadR*). Cells grown without supplement represent the calibrator condition. Error bars represent the standard deviations based on at least three independent biological replicates with duplicate qPCR reactions (NS, *P* >0.05; ***, *P* <0.005).

### Role of FabR/FadR in response to exogenous fatty acids

Given the strong exogenous fatty acid phenotype, it was suspected that either FadR and/or FabR, which are known to regulate *E. coli fabA* and *fabB* in response to exogenous fatty acids, may be responsible for this regulatory phenomenon. Homologs of *fabR* (locus tag: PBPRA3467) and *fadR* (locus tag: PBPRA2608) were readily identified in the SS9 genome via homology searches and deletion mutants of both genes were generated in both SS9R and *pfaA∷lacZY* strain backgrounds (Table 1). As shown in Figure 3A, strains containing the Δ*fadR* mutation displayed decreased β-galactosidase activities. Comparing β-galactosidase activities of the *pfaA∷lacZY* reporter strains containing either *ΔfadR* or *ΔfabRΔfadR* deletions indicated that both strains have similar β-galactosidase activities and further ruled out the involvement of FabR in the regulation of the *pfa* operon. Interestingly, β-galactosidase assays of the *ΔfabRΔfadR* mutant indicated that the down regulation of the *pfaA∷lacZY* gene fusion in the presence of Tween 80 (18:1) was independent of both FadR and FabR (Figure 3A). Transcript abundance analyses performed on RNA samples from the *ΔfabRΔfadR* mutant grown in the presence or absence of Tween 80 (18:1) indicated that both *pfaA* and *pfaD* transcripts were down regulated in response to supplementation while *fabA* and *fabB* transcripts were essentially equivalent between the two conditions (Figure 3B). Fatty acid analyses of the corresponding single and double mutant derivatives of SS9R are shown in Supplemental Table 2 and indicate that strains containing the *ΔfadR* mutation had decreased EPA levels consistent with the reduced β-galactosidase activities noted. As predicted, given the role of FadR as a positive regulator of *fabA/B*, a decrease in monounsaturated fatty acids was noted in strains containing the *ΔfadR* mutation (Supplemental Table 2).

**Table 1.**
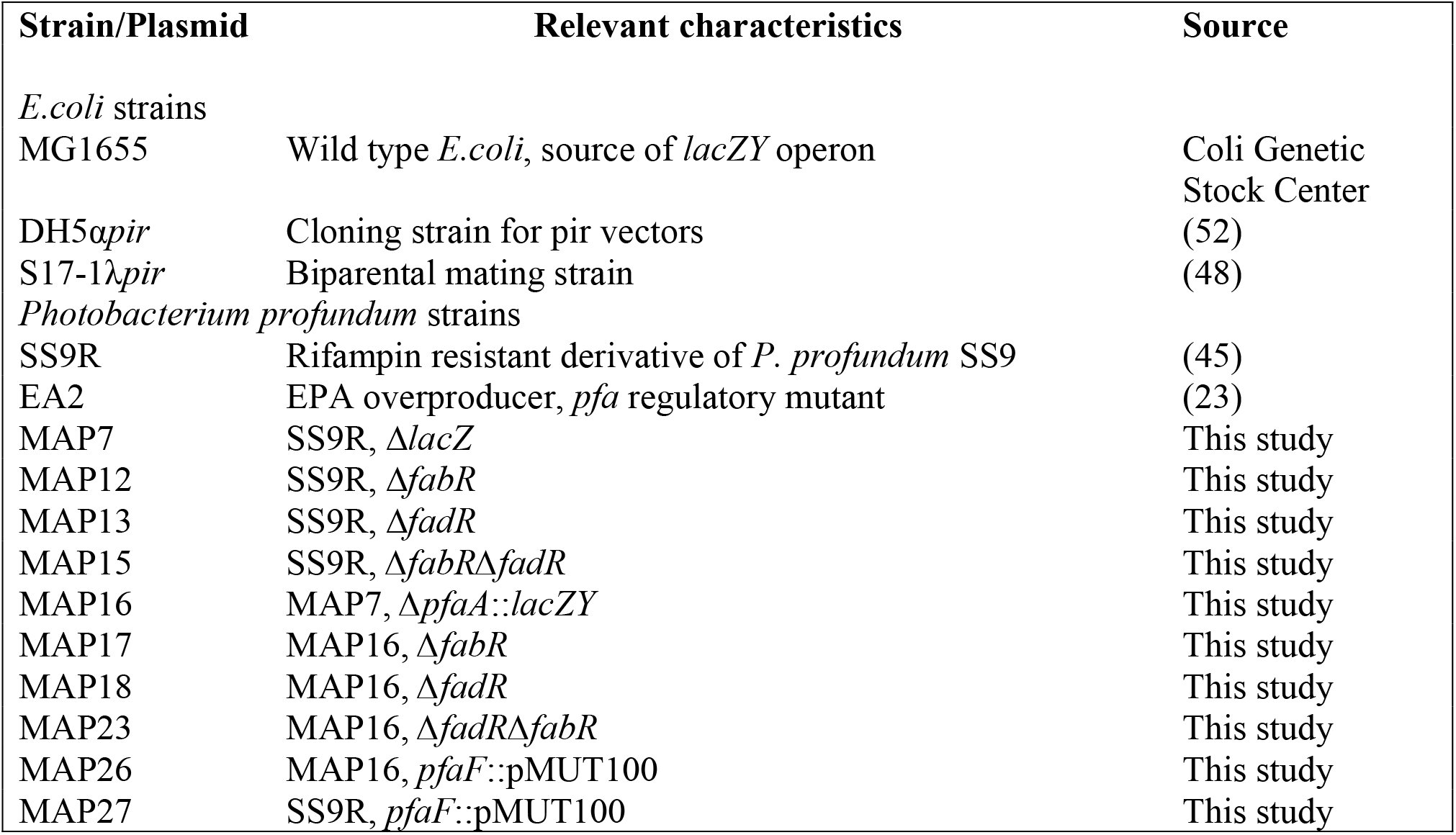
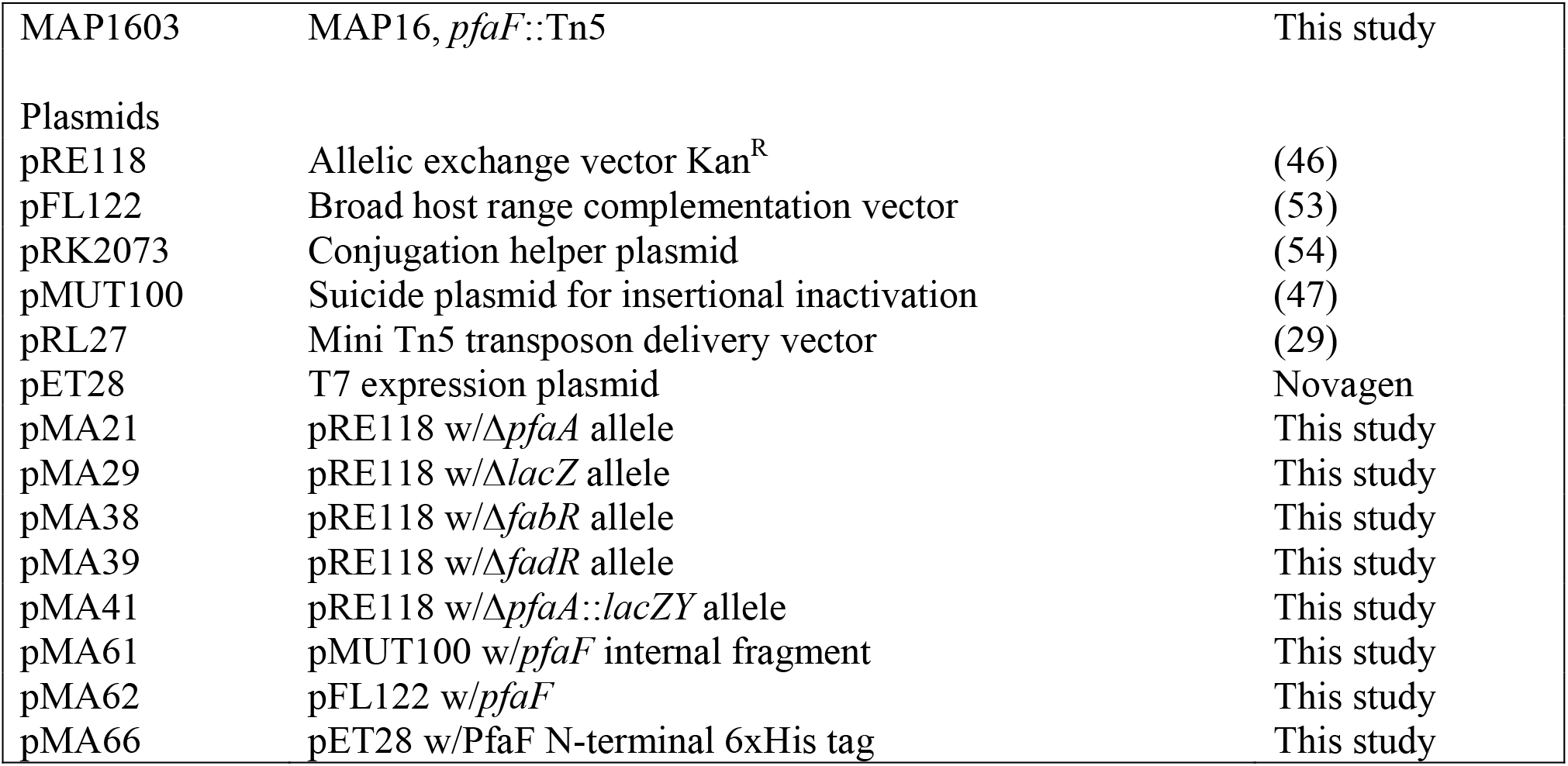
Strains and Plasmids used in this study

**Table 2.** Fatty acid compositions of SS9R and various *pfaF* mutant strains at 15°C

| Mean % fatty acid by weight<sup>a</sup> |  |  |  |  |
| --- | --- | --- | --- | --- |
| Fatty acid | SS9R | *pfaF* | *pfaF* pFL122 | *pfaF* pMA62 |
| **12:0** | 4.16 ± 1.53 | 2.84 ± 0.33 | 3.72 ± 0.45 | 3.73 ± 0.10 |
| **14:0** | 4.33 ± 1.21 | 5.25 ± 0.94 | 9.46 ± 0.31 | 8.03 ± 0.22 |
| **14:1** | 3.19 ± 1.11 | 3.90 ± 0.73 | 6.04 ± 0.19 | 4.90 ± 0.16 |
| **16:0** | 21.07 ± 1.84 | 19.03 ± 0.67 | 19.39 ± 3.64 | 19.50 ± 2.82 |
| **16:1** | 45.09 ± 1.68 | 50.95 ± 0.53 | 45.33 ± 2.06 | 39.59 ± 1.58 |
| **12-OH** | 1.99 ± 1.23 | 2.22 ± 0.28 | 2.66 ± 0.14 | 2.96 ± 0.27 |
| **18:0** | 0.71 ± 0.11 | 0.94 ± 0.08 | 1.37 ± 0.15 | 1.74 ± 0.09 |
| **18:1** | 12.26 ± 2.26 | 12.92 ± 0.42 | 9.38 ± 0.97 | 12.02 ± 2.82 |
| **20:5** | 4.52 ± 1.55 | 0.82 ± 0.23 | 1.14 ± 0.17 | 5.14 ± 0.30 |

<sup>a</sup> Data represents the average of at least three independent cultures ± standard deviation

### Identification of a regulator specific to the *pfa* operon

The FadR/FabR independent down regulation of the *pfa* operon in response to Tween 80 (18:1) observed here suggested that another regulator(s) for the *pfa* operon might exist in *P. profundum* SS9. To search for additional regulators, the *pfaA∷lacZY* strain was subjected to transposon mutagenesis using the mini-Tn5 delivery vector pRL27(29, 30). Of the approximately 10,000 mutants screened, several LacZ down or loss of LacZ activity mutants were identified and saved for further analysis. Arbitrary PCR was performed to identify the sites of mini-Tn5 insertion in these mutants. Excluding mutants that had Tn5 insertions in the reporter gene and/or *fadR*, we identified mutants from independent libraries that contained unique transposon insertions in the same gene PBPRA0221 (TetR bacterial transcriptional regulator, pfam13972) (Figure 4A). One of these mutants displayed a five-fold decrease in LacZ activity and no longer responded to exogenous Tween 80 (18:1) supplementation (Figure 4B). The PBPRA0221 gene, herein designated *pfaF*, encodes for a protein that is a member of the TetR family of transcriptional regulators and is clustered with genes related to lipopolysaccharide synthesis. To verify that this locus is involved in *pfa* gene regulation, single crossover insertion mutants were generated in the parental *pfaA∷lacZY* reporter strain and the EPA-producing SS9R strain. The LacZ activity of the resulting strain was identical to that seen in the *pfaF∷*Tn5 mutant thereby verifying this relationship and excluding the possibility of additional transposon insertion events being responsible for the observed phenotype (data not shown). Transcript abundance analysis of the *pfaF* mutant strain indicated significant down-regulation of both *pfaA* and *pfaD* transcripts relative to SS9R (Figure 4C) consistent with the LacZ activity data. Additionally, quantification of *fabA* and *fabB* transcripts indicated no major differences between SS9R and its *pfaF* mutant derivative. A comparison of the fatty acid profiles of SS9R and a *pfaF* mutant grown at 15°C shows the mutant displays an approximate 4-fold reduction in EPA content relative to the wild-type (Figure 3). The minor difference in the abundances of other fatty acids in the *pfaF* mutant is also consistent with the minimal changes in *fabA* or *fabB* transcription. In the case of both SS9R (EPA^+^) and *pfaA∷lacZY* (EPA^−^) strains, genetic disruption of *pfaF* did not lead to alteration in growth capabilities at high pressure or cold temperatures (data not shown).

**Figure 4.**
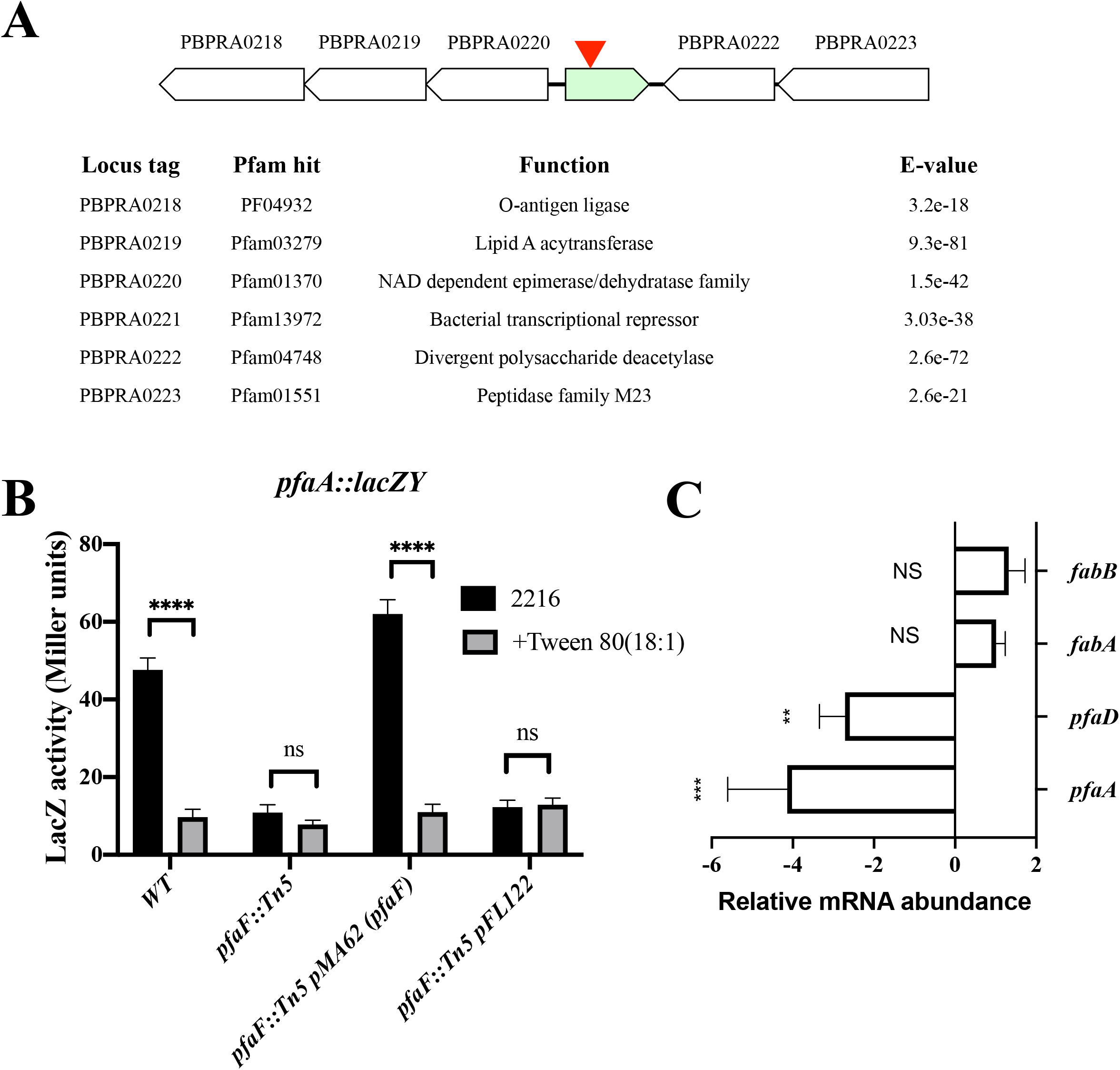
The *pfaF* genetic locus and regulatory phenotypes associated with its disruption. (A) Genetic organization of PfaF locus, arrow indicates relative position of mini-Tn5 insertion site in the *pfaA∷lacZY* regulatory mutant. (B) LacZ activity of *pfaF∷*Tn5 mutant and the complemented strains carrying pMA62 (pFL122 *pfaF*) or vector (pFL122) only ± 0.05% Tween 80 (18:1) supplementation (ns, *P* >0.05; ****, *P <*0.0001). (C) qRT-PCR analysis of *pfaA, pfaD, fabA*, and *fabB* transcript abundances in SS9R *pfaF* mutant relative to SS9R. Error bars represent the standard deviations based on at least three independent biological replicates with duplicate qPCR reactions (ns, *P >*0.05; **, *P* <0.005; ***, *P* <0.001).

The *pfaF* gene was cloned onto the broad host range complementation plasmid pFL122 and the resulting construct (pMA62) was introduced into *pfaF∷*Tn5 and SS9R *pfaF* mutant strains. As shown in Figure 4B, expression of *pfaF* was able to fully restore LacZ activities to that seen the parental *pfaA∷lacZ* strain relative to the vector only control. Similarly, the ability to down regulate the operon in response to exogenous Tween 80 (18:1) was restored in the complemented strain and not in the vector only control (Figure 4B). Fatty acid analysis of the complemented strains indicated that expression of *pfaF* from this construct was able to fully restore EPA levels back to wild-type levels compared to the vector only control (Table 3). Similarly, transcript abundances of *pfaA* and *pfaD* were restored to wild-type levels in the complemented strain relative to vector only controls (data not shown).

### Characterization of PfaF binding to *pfa* promoter

To verify that regulation by PfaF was direct, electrophoretic mobility shift assays (EMSA) were conducted. PfaF was cloned onto pET28 with an N-terminal 6x-His-tag to yield the construct pMA66 (Table 1). Using this construct, PfaF was expressed in *E. coli* and purified to homogeneity by Ni-affinity chromatography (Supplementary Figure 1). The DNA probe used in the EMSA was generated by PCR using a 6-carboxyfluorescein (6-FAM) labeled primer set and was designed to cover 400 base pairs directly upstream of the translational start of PfaA. As shown in Figure 5A, recombinant PfaF was able bind to the *pfaA* promoter probe in a concentration dependent manner forming a single discrete complex. Inclusion of a non-FAM labeled competitor probe at approximately 80-fold molar excess relative to the FAM labeled probe partially reversed the binding indicating that the interaction between PfaF and the probe is specific (Figure 5B).

**Figure 5.**
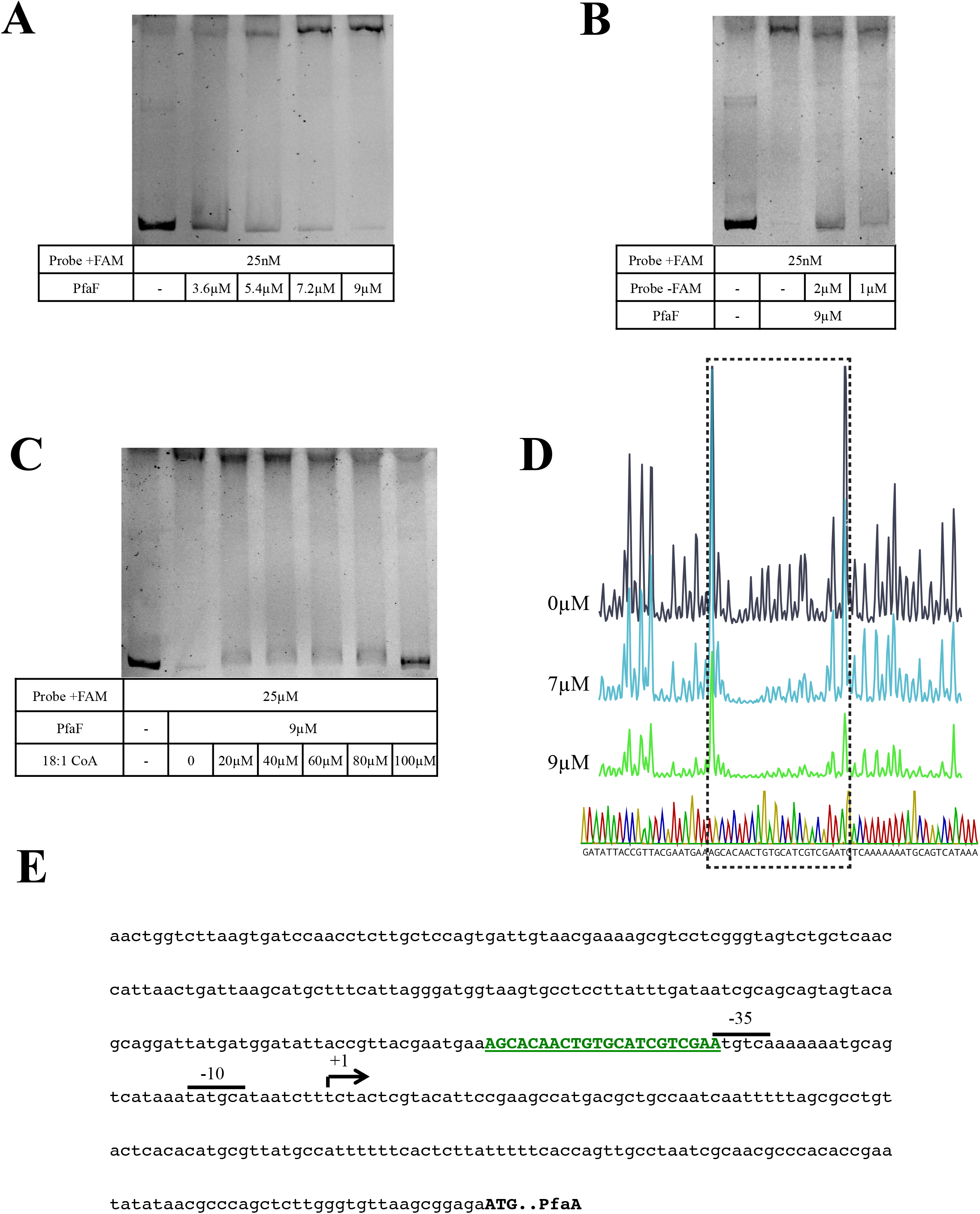
Characterization of PfaF binding to the *pfaA* promoter. (A) Electrophoretic mobility shift assay demonstrating PfaF binding to the *pfaA* promoter in a concentration dependent manner. (B) Binding of FAM-labeled probe (+FAM) is partially inhibited by the inclusion of molar excess of unlabeled probe (-FAM) indicating that PfaF binding is specific. (C) Addition of oleoyl-CoA at the indicated concentrations reverses the binding of PfaF to the probe in a concentration dependent manner. (D) DNase I footprinting analysis of PfaF binding to *pfaA* promoter. Purified PfaF was added at the indicated concentrations and subjected to DNase I digestion as described in Materials and Methods. Chromatograms and sequencing traces shown correspond to the coding strand and the box indicates the region protected from digestion by PfaF. (E) DNA sequence of *pfaA* probe used in mobility shift and footprinting assays. Putative promoter elements (−35 and −10 sites) and transcriptional start site (arrow) were previously determined (25). Region protected by PfaF indicated in bold, green, underlined font.

Further phylogenetic analyses of PfaF indicated that homologs exist in other polyunsaturated fatty acid producing bacteria such as members of the genus *Shewanella* and *Colwellia*, and that these homologs are distinct from other well known transcriptional regulators associated with fatty acid biosynthesis (Supplemental Figure 2A). Further investigation showed that that one such homolog from *Shewanella amazonensis* SB2B (PBD: 3rh2) had an unpublished crystal structure (Supplemental Figure 2B). Analysis of this crystal structure of the *Shewanella* PfaF homolog indicated that its C-terminal domain contained a ligand pocket containing an unknown ligand resembling the hydrocarbon “tail” of a fatty acid (Supplemental Figure 2C). Given the *in vivo* responses to the various fatty acid supplements observed, it was suspected that PfaF binding activity could be mediated by oleoyl-CoA (18:1-CoA). Mobility shift assays indicated that the addition of oleoyl-CoA abolished PfaF binding activity in a concentration dependent manner (Figure 5C).

To localize the binding site for PfaF in the *pfaA* promoter, non-radioactive DNase I footprinting assays were conducted utilizing the 6-FAM labeled probe used in the EMSA analysis. As shown in Figure 5D, inclusion of increasing amounts of purified PfaF led to protection of a 22 base pair sequence (AGCACAACTGTGCATCGTCGAA) corresponding to positions −182 to −204 of the promoter probe. The identified binding site is positioned directly upstream of the previously identified −35 site within the *pfaA* promoter (Figure 5E).

## Discussion

In this work, the genetic regulation of the *pfa* operon has been extensively characterized utilizing a variety of genetic techniques. While the omega-3 polyunsaturated fatty acid products (13, 14), biosynthetic mechanism (15, 18, 31), and phylogenetic distribution (16, 32) of the bacterial *pfa* operon has been extensively studied, there has been little work done describing how the operon is regulated and what gene(s) might be involved. The findings described in this report represent the first systematic investigation into the genetic regulation of the *pfa* operon. The finding that neither high hydrostatic pressure nor low temperature affected the activity of the *pfaA∷lacZY* reporter is consistent with previously reported results in *Photobacterium profundum* SS9 (25) and validated our reporter gene fusion approach. As noted previously in *P. profundum* SS9 (25), we confirmed the lack of correlation between the expression level of the *pfa* operon and the proportion of EPA found in the membrane phospholipids under cold temperature and high pressure culture conditions. The reasons for this phenomenon are unclear but they suggest that other factors in the biosynthesis and membrane incorporation of EPA are involved in the increased abundance of EPA at high pressure and/or cold temperature.

The finding that the *pfa* operon was down regulated in response to exogenous fatty acids in a FadR/FabR independent manner indicated that another transcription factor was responsible for regulating this response to fatty acid supplementation. Screening of a transposon library in the *pfaA∷lacZY* reporter strain identified a novel regulator, *pfaF*, whose gene product acts as a positive regulator of the *pfa* operon. Reintroduction of a null mutation in *pfaF* in SS9R yielded a mutant that had a specific several fold decrease in EPA composition with relatively minor changes in the abundances of the other fatty acids in the membrane. Successful complementation *in trans* confirmed the role of *pfaF* in positive regulation of the *pfa* operon. Based on amino acid sequence, PfaF is a member of the TetR transcriptional regulator family, of which several members have been characterized to be involved with regulation of fatty acid biosynthesis and/or degradation in other bacteria (33).

Mobility shift and DNase I footprinting analyses verified that PfaF is capable of binding to the *pfaA* promoter and localized its binding site within the *pfaA* promoter. The binding site of PfaF and its position relative to the transcriptional start of *pfaA* is typical of other positive transcriptional regulators (34) and further supports the role of PfaF as the transcriptional activator of the *pfa* operon. The *in vivo* data indicated that PfaF regulates the *pfa* operon in response to exogenous fatty acid supplementation, with the most robust response mediated by a long-chain monounsaturated fatty acid (18:1). As expected, addition of oleoyl-CoA disrupted PfaF binding activity in a concentration dependent manner. Regulation of the operon in both SS9R and *pfaA∷lacZY* strains by PfaF also indicates that the final EPA omega-3 fatty acid product of the Pfa synthase is most likely not involved with mediating this regulatory activity. While the physiologically relevant ligand(s) of PfaF has not been determined, the data presented in this work strongly suggests that it is a fatty acid or a derivative thereof, e.g. acyl-CoA or acyl-ACP.

Based on these results, a proposed model of regulation is shown in Figure 6. In the absence of fatty acids, PfaF binds to its cognate sequence within the *pfaA* promoter region and acts as a positive regulator. In the presence of exogenous fatty acids, which are presumably converted into acyl-CoA, PfaF binding to an acyl-CoA leads to its dissociation from the promoter leading to a lack of transcriptional activation of the *pfa* operon. Given its role in producing fatty acids for phospholipid biosynthesis (14) and its utilization of the same precursor metabolites as the Type II fatty acid synthase (15), it is not surprising that the Pfa synthase is controlled in a nearly identical fashion, albeit with its own cognate regulatory protein.

**Figure 6.**
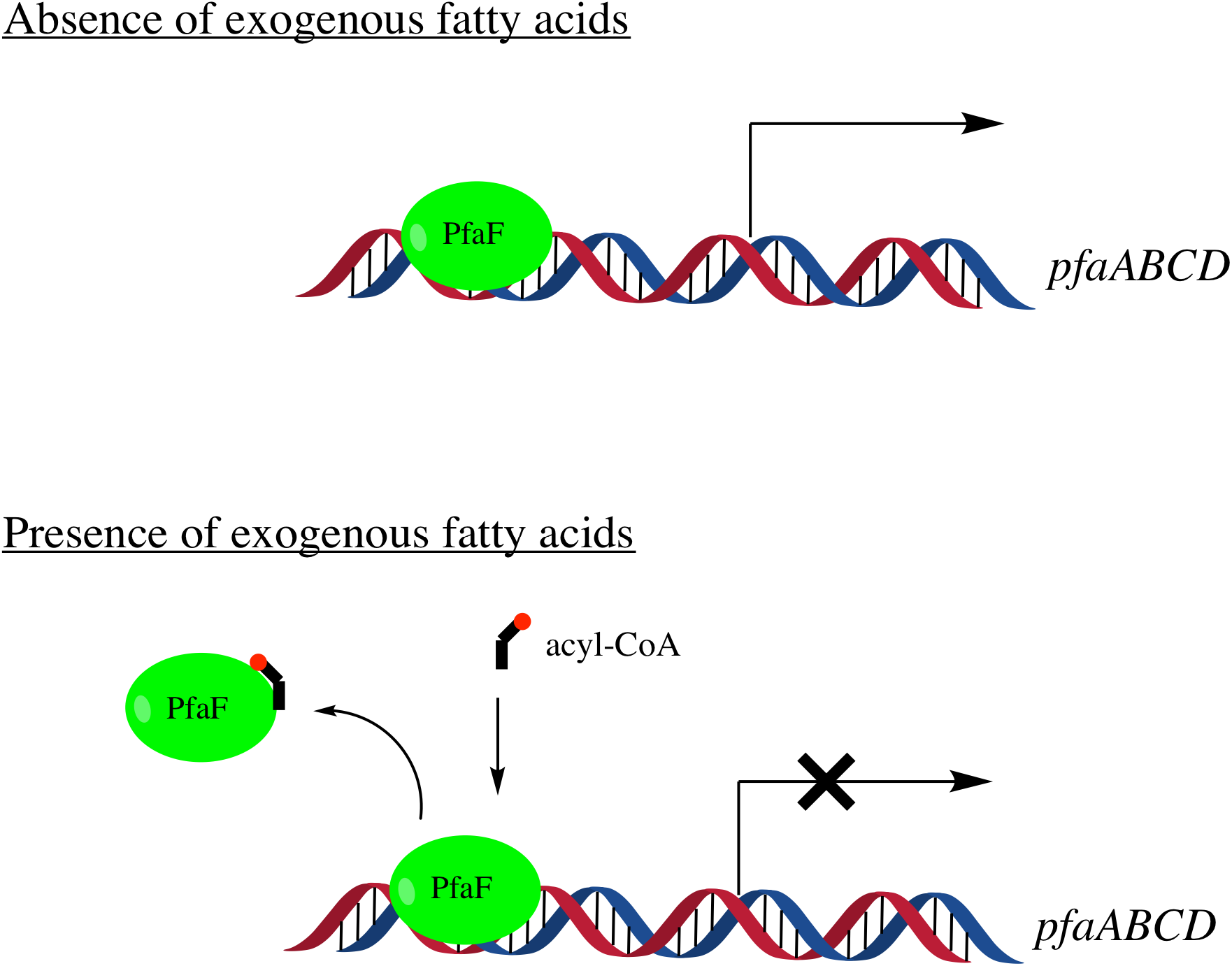
Proposed model of regulation of the *pfa* operon as mediated by PfaF. In the absence of fatty acids PfaF binds to the *pfaA* promoter and acts as a positive regulator. In the presence of exogenous fatty acids PfaF binds to acyl-CoA and releases from the promoter region leading to down regulation of the *pfa* operon.

Interestingly, genes regulated in a similar fashion, typically by FadR, have been shown to be involved in processes related to virulence amongst various members of the Vibrionaceae (35–38). Furthermore, many of these virulence genes are regulated in response to exogenous fatty acids (39), similar to the results reported here for the *pfa* operon. Whether the production of EPA or other PUFA is part of a virulence, symbiosis, or colonization response is unclear, although it is interesting to note that many PUFA-producing bacteria have been isolated from the guts, glands, or other tissues of marine animals such as fish and various invertebrates (40, 41).

At this time it is unclear as to how the various Tween derivatives and their fatty acid components are utilized by *P. profundum* SS9. Various members of the Vibrionaceae genus have been shown to have outer membrane associated lipase enzymes capable of hydrolyzing esterified fatty acids for incorporation into phospholipids or usage as a carbon source via β-oxidation (42, 43). In particular one such enzyme designated VolA, has been well-characterized as being involved with utilization of exogenous esterified fatty acids such as lysophosphotidylcholine (42). Preliminary searches of the *P. profundum* SS9 genome located a homolog (locus tag: PBPRA2574) of *volA* in an operon with a FadL homolog, identical to the genomic context of *volA* in *Vibrio cholerae.* This lipase or other similar lipases may be involved with the utilization of Tween compounds as a lipid source or as a carbon source.

Homology searches of available genomes indicated that all PUFA-producing marine Gammaproteobacteria contain a *pfaF* homolog many of which residing in similar genomic contexts. Whether *pfaF* is involved with regulating the *pfa* operon in these strains has yet to be determined. In many instances, such as in strains of *Shewanella* and *Colwellia*, the *pfa* operon also contains an annotated regulator typically designated as *pfaR*, immediately upstream of *pfaA.* Interestingly, the protein sequence of this regulator does not match to any class of bacterial transcriptional regulators and only contains an identifiable N-terminal helix-turn-helix domain, which is most likely involved in DNA-binding activity. Preliminary results using *S. piezotolerans* WP3, a genetically tractable EPA producer, indicated no differences in EPA composition between wild-type and *ΔpfaR* mutants under a variety of temperatures (data not shown). A previous study (44) demonstrated that replacing *pfaR* with an inducible promoter could lead to dramatic increases in EPA production in a heterologous host strain of *E.coli.* Unfortunately, that study lacked adequate data that could be used to ascertain the role of *pfaR* directly. Regulation of the *pfa* operon in strains with *pfaR* may indeed be more complex or otherwise different than in the case of *P. profundum* SS9.

The results presented here describe the identification of a novel transcriptional regulator in the model marine bacterium *P. profundum* SS9 that specifically modulates expression of the *pfa* operon in response to exogenous fatty acids and controls the amount of polyunsaturated fatty acid incorporated into membrane phospholipids. This study adds new insight into the unique lipid physiology of widespread marine bacteria and offers new opportunities for the genetic optimization of microbial omega-3 polyunsaturated fatty acid synthesis.

## Materials and Methods

### Bacterial strains and growth conditions

*Escherichia coli* strains were routinely grown at 37°C in Luria Bertani (LB) media unless stated otherwise. *Photobacterium profundum* SS9 strains were grown at 15°C in 2216 marine broth (Difco) at 75% strength (28g/L) unless noted otherwise. For solid medias, agar was included at 15g/L. The antibiotics kanamycin (50 μg/ml for *E. coli* and 200 μg/ml for *P. profundum*), chloramphenicol (15μg/ml), ampicillin (100μg/ml), and rifampicin (100μg/ml) were used as required. For high-pressure growth experiments, *P. profundum* SS9 strains were grown in heat-sealed bulbs and incubated in stainless steel pressure vessels as described previously (45).

### Targeted Mutagenesis

Vectors for introducing mutations into *P. profundum* were introduced by conjugation using previously described methods with minor alterations (14, 45). In-frame deletions were generated by allelic exchange using the suicide vector pRE118 (46). Insertional inactivation of target genes was accomplished by introduction of the suicide plasmid pMUT100 as described previously (23, 47).

### Transposon mutagenesis and screening

Biparental conjugations using *E.coli* S17-1λ*pir* were used to transfer the mini-Tn5 delivery plasmid pRL27 into the *pfaA∷lacZY* reporter strain (Table 1) (29, 48). Both recipient and donor strains were grown to stationary phase and conjugations were performed as described above. After ~24hr at ambient temperature (~22°C) cells on filter membranes were resuspended in 2216 broth and plated onto selection media (2216 agar containing 200μg/ml kanamycin and 100μg/ml rifampin) and incubated at 15°C for 5 days. Resulting ex-conjugants were patched to fresh selection plates in grid format. After two days growth at 15°C the arrayed mutants were replica plated to 2216 agar with X-gal (80μg/ml). After two days of growth on indicator media, mutants were screened by eye for differences in blue colony formation. Mutants with differential *lacZ* activity were clonally isolated and further screened by β-galactosidase assays in liquid cultures as described below.

### Identification of transposon insertion sites

To identify transposon insertion sites of interest an arbitrary PCR method was utilized similar to the method described previously (30). Primer sequences are given in Supplementary Table 1. In the first round of PCR, a primer specific to one end of the mini Tn5 element (Tn5 ext) in combination with one of three degenerate primers (arb1, 2, or 3) is used with purified genomic DNA as a template. The conditions used for the first PCR were; 95°C 5 min, 6 cycles of 95°C for 30 sec, 30°C for 30 sec, 68°C for 2 min, followed by 30 cycles of 95°C for 30 sec, 45°C for 30 sec, 68°C 2 min, and 68°C 5 min. Two microliters of the first reaction was used as a template for a nested PCR with “Arb clamp” and “Tn5 int” primers. The conditions for the second PCR were 95°C for 5 min, followed by 30 cycles of 95°C for 30 sec, 55°C for 30 sec, 68°C for 2 min, followed by 68°C for 10min. PCR reactions yielding single amplicons as judged by agarose gel electrophoresis were purified using a PCR clean up kit (Zymo Research) and sent for DNA sequencing. Sequences were compared to the genome of *P. profundum* SS9 to determine the insertion site of the mini-Tn5 element.

### β-galactosidase assays

Cultures of indicated strains were grown at 15°C in aerobic tubes, unless noted otherwise. Mid-log phase cultures (OD_600_ = 0.2-0.6) were assayed for changes in LacZ activity from whole cell extracts using the SDS and chloroform lysis modification described previously (49). Activities reported are in Miller units and represent the mean of at least five independent experiments.

### RNA isolation and quantitative reverse transcriptase PCR (qRT-PCR)

Total RNA was isolated from mid-log phase cells grown under the indicated conditions using Trizol (Invitrogen) following manufacturer guidelines. Crude RNA extracts were further purified and treated with DNase I (Zymo Research) using the RNA Clean and Concentrator kit (Zymo Research). For cDNA synthesis, the Superscript III First Strand synthesis kit (Invitrogen) was used following manufacturer’s recommended protocols. Quantitative PCR’s were performed using the Maxima Sybr Green Master Mix (Thermo Scientific) and run on a Stratagene MX3000P qPCR system. For quantification of target transcripts, the *gyrB* gene (PBPRA0011) was used as an internal reference and differences in expression were calculated using the ΔΔCT method. Primers for qPCR experiments are listed in Supplementary Table 1.

### Expression and Purification of PfaF

PfaF (locus tag: PBPRA0221) was cloned into pET28 as an NheI-XhoI fragment (Novagen) as to generate N-terminal 6xHis tagged protein. After sequence verification, this construct was transformed into BL21 DE3 Tuner pLysS (Novagen) cells following standard procedures (49). For protein expression, overnight cultures were diluted 1/100 into LB supplemented with chloramphenicol (30μg/ml) and kanamycin (50μg/ml) and grown at 30°C until OD_600_ of ~0.5 at which point IPTG was added to final concentration of 0.5mM and grown for an additional 4hrs at 30°C. Cells were harvested by centrifugation and cell pellets were processed or stored at −80°C. Frozen cell pellets were thawed on ice with the addition of buffer A (50mM Tris-Cl pH 7.5, 200mM NaCl, 10% glycerol). Lysozyme was added and cells incubated on ice for 30 minutes and sonicated on ice to complete the lysis procedure. The lysate was centrifuged at 10,000 rpm, 4°C, for 30 minutes to separate insoluble and soluble fractions. The clarified supernatant was applied to a Ni-NTA column equilibrated with buffer A and mixed gently at 4°C for one hour. The resin was washed with several column volumes of buffer B (buffer A + 30mM imidazole) and proteins were eluted with buffer C (buffer A +300mM imidazole). Eluted fractions were checked by SDS-PAGE for purity, and appropriate fractions were pooled and desalted using PD-10 columns (GE Healthcare) and exchanged into Buffer D (20mM Tris Cl pH 7.5, 50mM NaCl, 10% Glycerol). Protein samples were pooled and subsequently concentrated by 10,000 kDa centrifugal filter units (Amicon).

### Electrophoretic mobility shift assays and DNase I footprinting

A 400bp DNA fragment, which included the previously mapped promoter of *pfaA*, was generated by PCR using a 6-carboxyfluorescein (FAM) labeled primer listed in Supplemental Table 1 and purified *P. profundum* SS9 genomic DNA as a template. For mobility shift experiments, binding reactions contained; binding buffer (20mM Tris-Cl pH 7.5, 0.2mg/ml BSA, 0.5mM CaCl_2_, 2.5mM MgCl_2_, 10% glycerol), 2μg poly dI-dC (Thermo Scientific), 25nM promoter probe, and the indicated amount of purified PfaF. Binding reactions were incubated at 22°C for 60 min and analyzed by electrophoresis using pre-run 6% 0.5X TBE, 1% glycerol, native polyacrylamide gels (50). Gels were visualized and photographed using a GelDoc system (Bio-Rad).

Non-radioactive DNase I footprinting (51) was performed using the same binding buffer and reaction conditions used in the EMSA experiments described above. Digests were initiated by the addition of 0.03U of DNase I (NEB Biolabs), and incubated at 22°C for 2 minutes. Digestion reactions were stopped by the addition of DNase I stop buffer (NEB) and were extracted with phenol:chloroform:isoamyl alcohol (25:24:1, Thermo Scientific). DNA fragments were further purified by use of a PCR purification kit (Zymo). Eluted DNA fragments were subjected to fragment analysis by capillary electrophoresis (Eton BioScience). Chromatograms were examined using the microsatellite plugin within Geneious Prime 2019.2.3 (https://www.geneious.com). Protected regions were identified and compared to the included LIZ-500 size standards to identify the binding site coordinates and relevant protected bases.

### Fatty acid extraction and GC-MS analysis

Late log phase cultures were harvested by centrifugation and cell pellets rinsed once with 50% Sigma Sea Salts solution (16g/L) and stored at −80°C. Cell pellets were lyophilized and fatty acids were converted to fatty acid methyl esters and analyzed by gas chromatography mass spectrometry using previously described protocols and methods (14).

## Acknowledgements

We would like to thank Dr. Doug Bartlett for insightful discussions regarding genetic manipulations in *P. profundum* SS9 and Dr. Bianca Brahamsha for the generous gift of the pRL27 mini-Tn5 transposon delivery vector. This work was supported by National Science Foundation Division of Molecular and Cellular Biosciences grant MCB-1149552 to E.E.A.

